# Drought drives reversible disengagement of root-mycorrhizal symbiosis

**DOI:** 10.1101/2025.08.25.671999

**Authors:** Garo Z. Akmakjian, Kazunari Nozue, Hokuto Nakayama, Alexander T. Borowsky, Angel M. Morris, Kristina Baker, Alexander Cantó-Pastor, Uta Paszkowski, Siobhan Brady, Neelima Sinha, Julia Bailey-Serres

## Abstract

The increasing frequency and severity of droughts pose a major threat to agriculture, food security and ecosystems. Plants respond to water deficit by adjusting growth and metabolism to enhance survival; these adjustments impact the soil microorganisms interacting with plant roots. Plants establish symbiotic relationships with arbuscular mycorrhizal fungi which supply soil nutrients in exchange for carbon metabolites via an intricate dual-species interface within roots. These fungi are dependent upon host-derived photosynthates and are thus potentially vulnerable to plant perturbations during drought. Here, we demonstrate that the plant-mycorrhizal relationship is dynamic when water becomes limiting. During water deficit, rice de-prioritizes nutrient acquisition gene regulatory networks, including its AM symbiotic program, in a strategy conserved with tomato. The fungal symbiont correspondingly represses its growth, undergoing metabolic quiescence, coupled with decommissioning of hyphae within the host’s root. Following re-watering, the host re-engages with its partner fungus, re-invigorating fungal growth and arbuscule establishment. This coordinate, reversible and enduring inter-organismal association may aid host survival under transient stress, but suggests that mutualisms in native and crop plants are potentially fragile in increasingly erratic climates.

## Main

Nearly 80% of land plants can establish symbiotic interactions with arbuscular mycorrhizal fungi (AMF), filamentous Glomeromycotina fungi that colonize root cortical cells with specialized branched hyphal structures called arbuscules. These tree-shaped dual-organism membrane interfaces mediate the efficient provision of soil-derived minerals captured by fungus to the plant in exchange for fixed carbon in the form of sugars and lipids. Nutrient transfer to the plant is predominantly of phosphate (Pi), but AM symbiosis purportedly confers broad resilience to environmental stressors, including drought^1–4^. It is unclear how transient stresses that disrupt the delicate economic balance of symbiosis influence the host-fungal interaction. Field studies suggest that AM symbiosis is inhibited as soil moisture declines in the dry-land crop sorghum^5^ whereas it may be induced as paddies dry in rice^6^, suggesting nuanced dynamic interactions.

Here, we investigated the inter-organismal dynamics of rice and the AMF *Rhizophagus irregularis* as soils dry and are rewatered. Our results illustrate a coordinated reduction in nutrient exchange during drought, marked by transitions in co-regulated inter-organismal gene networks. These expose a transition from fungal DNA synthesis and the cell cycle toward a quiescent state. This was accompanied by retirement of established intraradical hyphae and arbuscules. Reactivation of symbiosis upon re-watering demonstrates a symbiotic association that is highly responsive to rapid environmental perturbations.

## Results

### Water deficit dampens the symbiotic program

To better understand inter-organismal dynamics in a changing environment, we compared root and shoot crown (measured 1 cm from the shoot base) transcriptomes of rice continually well-watered (WW) or experiencing a water-deficit (WD) that significantly reduced leaf relative water content (Fig. S1). Samples were collected on the day that the leaves of WD plants rolled, a moderate non-lethal stress in our experimental system^7^. The plants had been cultivated with (Myc) or without (Non-myc) *R. irregularis* in Pi-depleted substrate in drained, aerobic pots for 6 weeks prior to the WD to allow robust fungal colonization (Fig. 1A, Fig. S1). The overall WD response was similar between Myc and Non-myc plants, including a strong and pronounced ABA response (Fig. 1B, cluster 5; Fig. S2B and Table S3). The clusters of WD down-regulated genes were enriched for genes associated with nutrient starvation, particularly Pi (cluster 1), iron (Fe) and zinc (Zn) (cluster 4). To evaluate whether this repression of nutrient uptake is a broad response to WD or specific to these aerobic, low-nutrient conditions, we compared the results to our previous WD dataset with plants grown in continuously-irrigated nutrient-replete conditions closely corresponding paddy-grown rice^7^. Nutrient starvation-associated genes were some of the most enriched amongst WD-repressed genes across experimental designs, including genes for Pi, Fe, Zn and nitrogen (Fig. S2). This suggests that regardless of nutritional status at the onset of WD, nutrient acquisition is heavily de-prioritized when water is limiting, perhaps as an energy saving mechanism.

**Figure 1.**
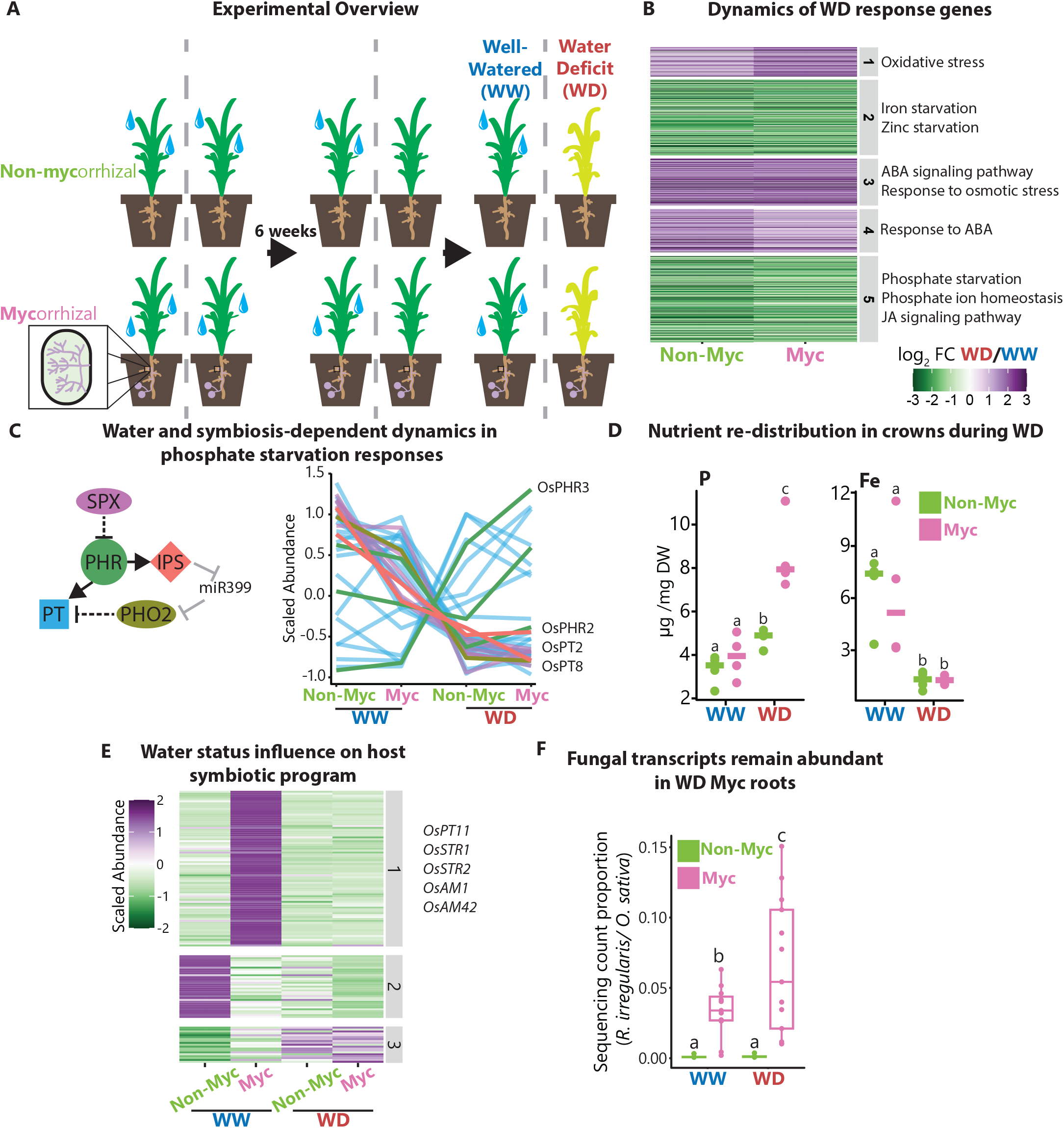
The mycorrhizal symbiotic program is repressed by water deficit. **(A)** Overview of experimental design. Colonized (Myc) and non-colonized (Non-Myc) rice plants were subjected to mild water deficit (WD). **(B)** Transcriptomic response of WD-response genes is similar between Myc and Non-Myc roots. WD DEGs (*p* < 0.05, |log_2_ FC| > 1) from either Myc or Non-myc roots were *k* means clustered. **(C)** The phosphate starvation response is repressed by WD. Dynamics of phosphate (Pi) starvation regulators and phosphate transporters (*OsPTs*) in response to AM and WD. (Left) Schematic of Pi deficiency signaling in rice. Each node represents a family of multiple genes. (Right) Line plots of Pi genes depicted in the schematic. Each line represents a separate gene and is color-coded according to the schematic. Computed *p* values determined by DESeq2 are available in Table S2. **(D)** Nutrients are redistributed in crowns during water deficit. ICP-OES analysis of Pi and Fe concentration in crowns (*n* = 4). Different letters indicate significantly different means based on Tukey’s test. Data for additional elements and replicates are in Fig. S4. **(E)** The host AM symbiotic program is repressed during WD. Transcriptomic response of the AM response module in Myc and Non-myc roots. **(F)** The host remains colonized during WD. Proportion of fungal-derived RNA-seq reads compared to rice-derived reads (*n* = 11-12). Different letters indicate significantly different means based on Tukey’s test.

Myc colonization was sufficient to repress genes associated with the Pi starvation response (PSR) under WW conditions (Fig. 1B and Fig. 1C), suggesting at least partial amelioration of Pi deficiency via symbiotic exchange. During WD, Pi uptake genes were broadly repressed regardless of colonization status, including genes that regulate the Pi deficiency response (*SPX1/2/3/4/5/6, IPS1/2, PHO2* and *PHR2*) as well as many Pi transporters in the *OsPT* family (Fig. 1C). Unlike mild WD which induces PSR genes^8^, more severe WD appears to repress PSR genes. However, the PSR transcriptional regulators *OsPHR3* and *OsPHR4* and a subset of *OsPT* transporter genes were *induced* by WD. Simultaneous corresponding changes in chromatin accessibility of *OsPHR2* and *OsPHR3* promoters also occurred during WD under Non-myc nutrient replete conditions (Fig. S3). The rice *PHR* homologs are thought to be partially redundant positive regulators of both PSR and AM symbiosis^9,10^. However, the differential regulation of *PHR* genes during WD suggests some level of sub-functionalization, potentially to regulate distribution of Pi. We observed an increase in crown P concentration following WD (Fig. 1D), particularly in the colonized plants. Concentrations of iron decreased drastically along with a modest increase in zinc (Fig. S4) concomitant with expression changes in genes encoding their transporters (Fig. S5). Given the repression of genes associated with cell division (Table S4, GO term table) and crown root initiation (e.g., *CROWN ROOTLESS1*), the function of enriched P in crowns during WD is likely not associated with active growth, but the reason is unclear.

The changes in nutrient uptake mRNAs in roots and elements in crowns, prompted evaluation of WD impact on AM symbiosis and plant nutrient acquisition. We found the majority of AM-inducible genes were repressed to non-Myc levels by WD (Fig. 1E). The abundance of transcripts canonically associated with symbiosis such as the GRAS transcription factor *OsRAM1* and putative fatty acid (FA) transporters *OsSTR1/2* were indistinguishable between WD Myc and WD Non-myc plants (Table S2). Symbiotic gene expression was repressed in a parallel water deficit experiment performed in tomato (Fig. S6), indicating broad conservation of this phenomenon in angiosperms. Signaling for the stress hormone ethylene has been shown to inhibit AM symbiosis via the karrikin-dependent negative regulator of symbiosis SMAX1^11^; however, symbiosis in *smax1* mutants was also repressed by WD, suggesting a mechanism independent of karrakin signaling (Fig. S7).

Despite the dynamics in host transcriptomic signature indicative of symbiosis, fungal RNA-seq reads were present in both WW and WD Myc root samples. Surprisingly, their abundance relative to host reads increased during WD (Fig. 1G). Despite the persistent physical colonization by the symbiont, there is an apparent dysfunction in symbiosis.

### AM fungi enter a state of quiescence during water deficit

The incongruity posed by the presence of AMF but lack of a symbiosis-associated transcriptome signature led us to explore the impact of WD on the fungal symbiont. To do so, we leveraged the recent high quality genome assembly for *R. irregularis*^*12*^ and complemented gene annotations with the construction of orthogroups based on 17 diverse fungal taxa to identify orthologues of genes from well-studied fungal species (Table S8). To facilitate direct comparisons of host and fungal transcripts, we mapped reads to both species simultaneously using a combined transcriptome. Of 30,209 annotated gene models, nearly 7,000 *R. irregularis* transcripts passed the detection threshold in our root tissue dataset.

We exploited the four-variable root transcriptome dataset to construct dual-species co-expression networks to identify fungal genes that are co-expressed with host marker genes. For example, a co-expression module containing major Pi (*OsPT11*^*13*^) and putative FA transporters (*OsSTR1/OsSTR2*^*14,15*^) and other core AM-inducible genes^16^ were found to be co-expressed with 580 fungal genes (Fig. 2A). The network includes nodes with high inter-species connectedness (Fig. S8). The most enriched GO terms for the fungal genes were related to chromatin structure and cell wall metabolism; indeed, multiple genes encoding each core histone subunit of nucleosomes were in this module (Fig. 2A). Binding sites for orthologs of *S. cerevisiae* transcription factors (TFs) that regulate cell cycle transitions^17–19^ were conserved and enriched in the promoters of these fungal genes (Fig. 2B, Table S9). An enrichment in binding sites for TFs promoting yeast growth in glucose but not non-fermentable carbon sources was found in the proximal 5’ regions of these genes^20–22^. The most strongly WD-repressed fungal genes included histones, which are indicators of mitotic DNA synthesis (S) phase activity (Fig. 2C). These observations support the hypothesis that both AM nuclear division and growth are associated with host propensity to maintain symbiosis and strongly inhibited by WD.

**Figure 2.**
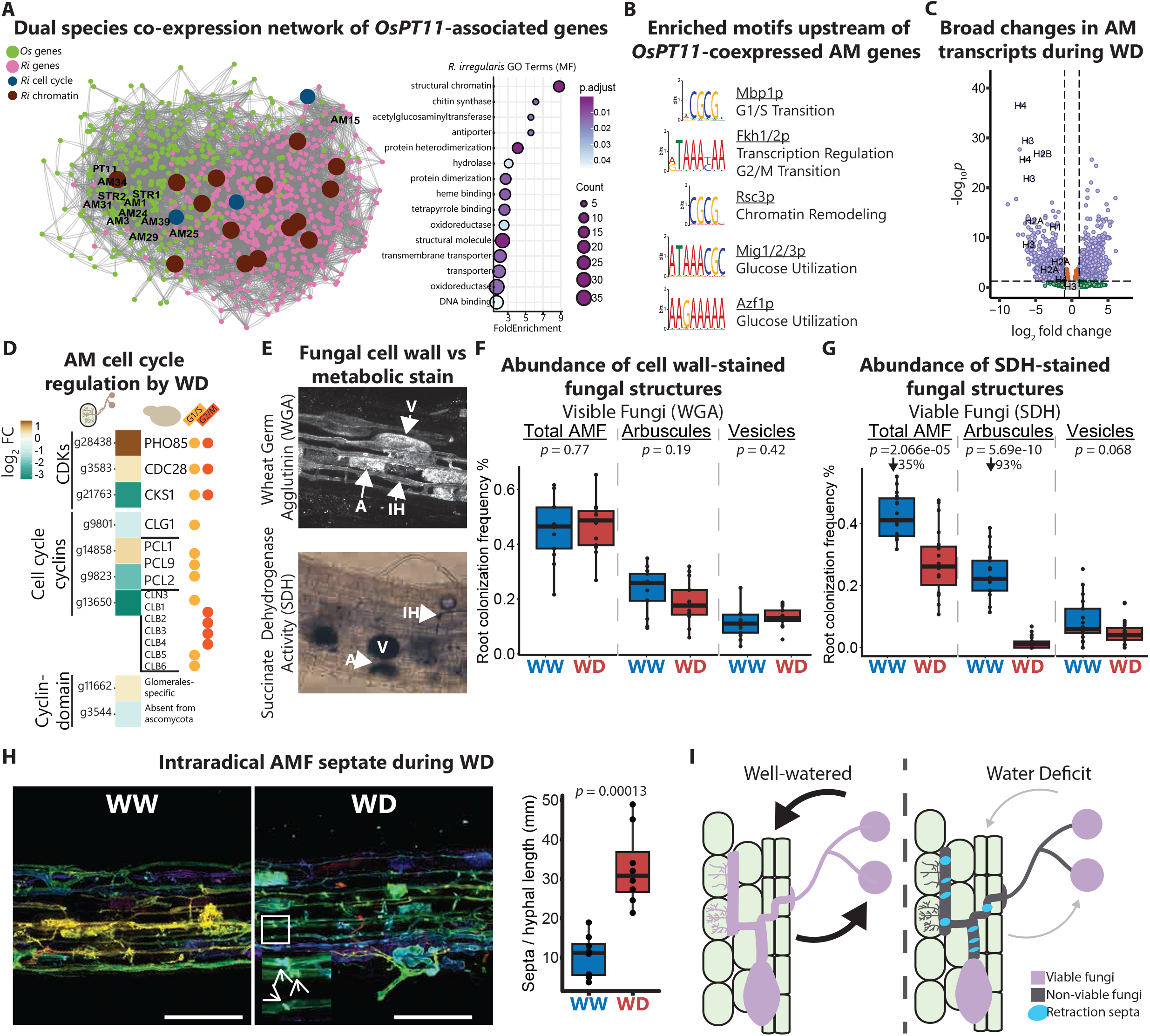
Intraradical AMF enter a quiescent state during drought. **(A)** Plant-fungal gene c-expression network resolve coordination of fungal growth with host symbiotic engagement. A multi-species network of genes co-expressed with host arbuscular mycorrhizal marker *OsPHOSPHATE TRANSPORTER11* (*PT11*). Symbiosis-associated rice genes and AM histone and cell cycle genes are highlighted. Right panel: Gene ontology group enrichment of fungal genes co-regulated with rice symbiosis genes. MF, molecular function. **(B)** Significantly enriched transcription factor binding motifs of AM genes co-expressed with rice symbiosis genes relate nuclear division and carbon use. **(C)** AM DNA synthesis genes are strongly repressed by WD. Volcano plot showing log_2_FC of fungal transcriptome in response to WD. Genes with |log_2_ FC| > 1 and *p* < 0.05 are shown in purple. **(D)** AMF repress the cell cycle during WD. Heatmap showing log_2_FC during WD of cyclin-dependent kinases, cell cycle-regulating cyclins and cyclin domain-encoding genes without clear orthology to well-characterized fungi based on OrthoFinder analysis. **(E-G)** Fungal structures remain intact during WD but are not metabolically active. **(E)** Representative micrographs of colonized roots stained with FITC-conjugated wheat germ agglutinin (WGA) forAM cell walls or succinate dehydrogenase (SDH) for viability. AM frequency per root length was assessed by the gridline intersect method for **(F)** WGA (*n* = 13) and **(G)** SDH (WW, *n* = 13; WD, *n* = 19). Different letters indicate significantly different means based on Tukey’s test. **(H)** Intraradical hyphae septate during WD. Colonized roots were stained with WGA and imaged by confocal microscopy. Left: Representative images of maximum projections of z-stacks for WW and WD roots. Right: Frequency of septa across hyphal lengths (*n =* 8). *p* value determined by Students *t*-test. **(I)**. Scheme summarizing plant-AM status in WW and WD states. Arrows indicate nutrient exchange.

To assess whether the fungus was actively undergoing nuclear division, we searched for genes associated with the cell cycle by identifying *R. irregularis* genes with cyclin-or cyclin-dependent kinase (CDK)-associated InterPro terms and followed by filtering for genes in orthogroups that contain cell cycle regulators of the yeasts *S. cerevisiae* or *Schizosaccharomyces pombe* (Fig. 2D). Levels of cyclin domain-encoding transcripts were largely repressed by WD. Interestingly, the number of cell cycle regulatory cyclins in *R. irregularis* and other AM species is significantly reduced compared to other fungi, particularly *S. cerevisiae*. An orthogroup of seven Cdc28-regulating cyclins in *S. cerevisiae* (CLB1-6 and CLN3) regulates different events at the G1/S and G2/M checkpoints but only one ortholog is recognizable in *R. irregularis* (*Ri*-*g13650*) and two other Glomeromycota (*Glomus cerebriforme* and *Gigantae rosea*) (Table S8). The singular ortholog of CDC28 was strongly repressed during WD. Phylogenetic analysis confirmed the lack of expansion of this cyclin subgroup in AM fungi compared to other eukaryotic lineages (Fig. S9). A similar lack of expansion was observed in *R. irregularis* for Pho85-regulating cyclins of the *S. cerevisiae* PCL gene family, which also included WD-repressed members (Fig. 2D), indicating Glomeromycota have reduced or altered checkpoint control. These data collectively suggest that the fungal symbiont arrests nuclear division during WD. We hypothesize that cell cycle arrest results in a quiescent state in AMF when water is limiting.

Given the transcriptomic evidence of growth cessation and symbiotic exchange (arbuscule) suppression, we ascertained whether fungal structures remained present during WD. In colonized roots stained with wheat germ agglutinin (WGA), which binds the fungal cell wall chitin, we found no difference in colonization frequency between WW and WD plants (Fig. 2E and 2F), based on assessment of intraradical hyphae, vesicles or arbuscules. There was a slight decrease in arbuscule frequency under WD, but this was not statistically significant. We next tested whether these fungal structures were metabolically active by staining for succinate dehydrogenase (SDH) activity, essential for mitochondrial electron transport. Transcripts encoding subunits of mitochondrial respiratory complex II, including the SDH catalytic subunits, were detectable in WD samples (Fig. S10), Nonetheless, there was a large decrease in SDH-stained AM fungi in WD plants, despite the maintenance of visible structures (Fig. 2E and 2G). Notably, we observed a near total loss of SDH-stained arbuscules during WD, whereas SDH-stained vesicles remained unchanged (Fig. 2G). This contrasting staining pattern suggests that while fungal presence is sustained during WD, the fungal symbiont arrests symbiosis and growth.

We questioned whether reduced SDH staining was indicative of inviability or simply curtailed metabolic activity as an energy conservation mechanism. *R. irregularis* are multinucleate fungi that do not septate to compartmentalize nuclei but form so-called retention septa that separate dead, hollow hyphal segments from viable regions. We quantified septation in WGA-stained roots as an indicator of death. To reduce potential bias due to variation in colonization maturity, we scored intraradical hyphae of root fragments that contain multiple fully developed arbuscules. Despite this requirement, septation was three times more frequent following WD (Fig. 2H, Video S1 and S2), indicating significantly more hyphal death. Consistent with the depletion of SDH-stained arbuscules, these results indicate that intraradical fungal structures become non-viable and that many WGA-stainable and SDH inactive fungal structures are AM “fossils” retained within roots (Fig. 2I).

### Colonization is re-established upon re-watering

The striking loss of viable arbuscules led us to investigate the recovery dynamics of the symbiosis. We repeated the water deficit (WD) experiment with AM-colonized plants but added a recovery (rewatering) period of up to 8 days (WDR), evaluating both shallow and deep roots in pots. One day post rewatering was adequate to repress the ABA response marker gene *OsRAB16a* to baseline levels in shallow and deep roots (Fig. 3A). Expression of the Pi deficiency marker *OsPT2* was restored to WW levels within this timeframe. This further supports the model that nutrient demand is acutely repressed during WD rather than alleviation of nutrient deficiency *per se*, and nutrient acquisition resumes when the threat of drought resolves, along with root growth^7^.

**Figure 3.**
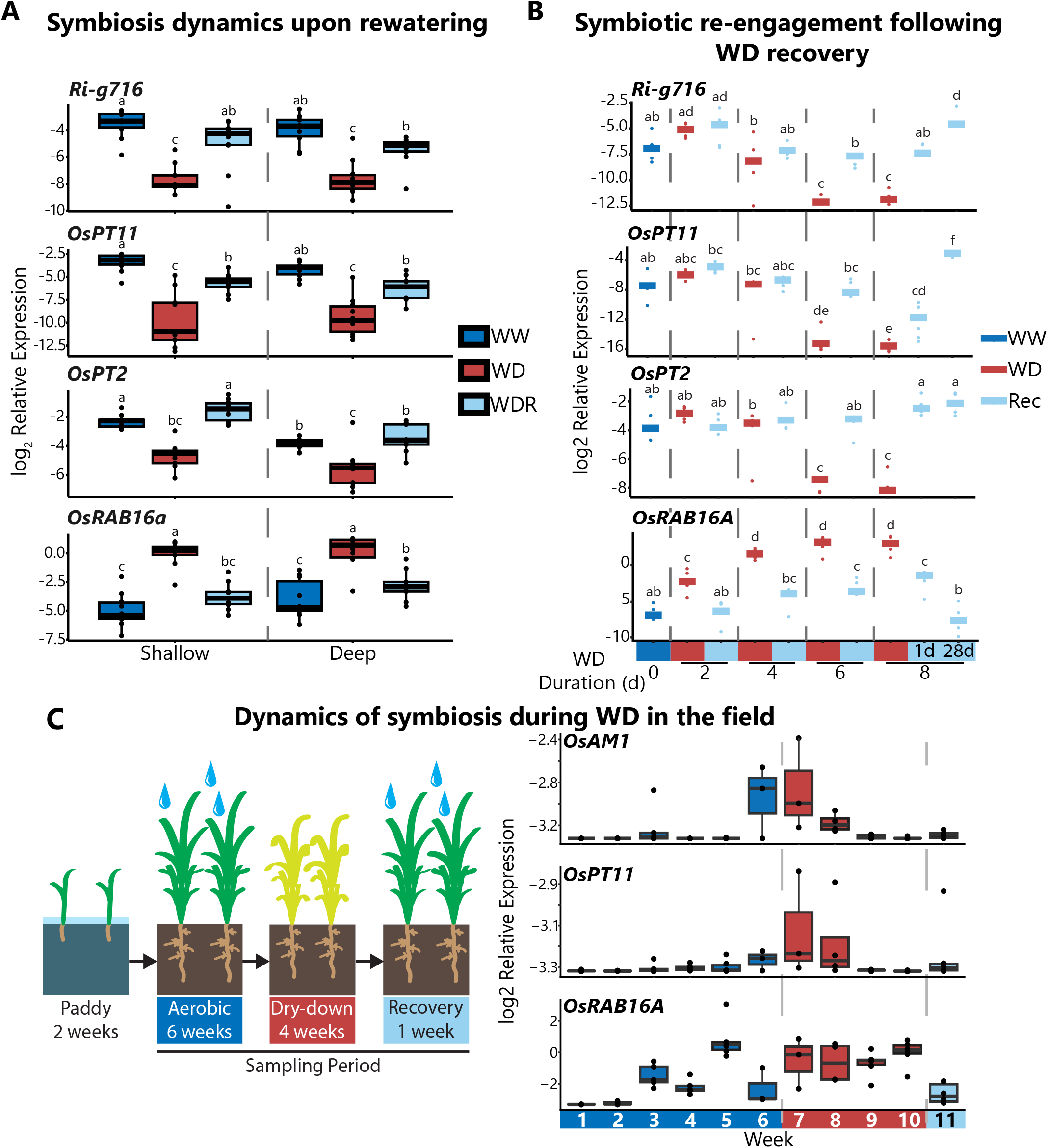
Symbiosis is re-invigorated during WD recovery (WDR). **(A)** Gene expression for S phase markers resumes upon re-watering but symbiosis is not immediately recovered. qPCR analysis of host and AM genes during WW, WD and WDR in roots separated into shallow (top 7.5 cm) and deep root compartments (7.5 cm) of pots. Plants were grown under WD until leaf rolling (5-6 d) and then re-watered for 24 hrs. Data are log_2_-transformed expression values, *n =* 9-10 pots. Different letters indicate significantly different means based on Tukey’s test within root compartments. **(B)** Arbuscule re-establishment is delayed after prolonged WD. Water was withheld for the indicated number of days, rewatered and sampled after 24 hr recovery; after 8 d WD an additional 28 d recovery time point was included. Data are log_2_-transformed expression values, *n =* 4-5. Different letters indicate significantly different means based on Tukey’s test across all time points. **(C)** AM symbiosis is drought responsive in the field. (Left) Schematic outline of field experiment. (Right) Gene expression of AM and drought markers genes (*n* = 4-6).

We also monitored transcripts associated with symbiosis. Levels of *Ri-g716* mRNA, which encodes a WD-repressed histone H3, rebounded to near WW levels during WDR. As a marker of S phase and thus nuclear division, this indicates that the *R. irregularis* rapidly breaks quiescence and likely resumes growth upon re-watering. On the other hand, *OsPT11* mRNA increased during WDR but was not fully restored to WW levels within 24 hr. Because water deficit progressively reduces moisture of the growth substrate and deeper roots are able to proliferate and extend lateral roots during WD and AMF preferentially colonize large lateral roots in rice, we hypothesized that deeper roots act as a reservoir of AMF. But the transcript changes were consistent between deep and shallow roots, suggesting that the reduction in colonization is systemic, at least in pot experiments. Collectively, these results show that symbiosis is transitory. While *R. irregularis* growth resumes upon re-watering, arbuscular symbiotic nutrient exchange is not maintained during WD and may not be restored upon re-irrigation, highlighting the dynamic interplay between the fungus and its host during transient environmental stressors such as WD.

To further evaluate temporal recovery dynamics, we explored how the duration, and thus severity, of WD impacts recovery of symbiosis. Plants were subjected to WD for 2, 4, 6 or 8 days with a 24 hr recovery for each WD duration. *OsRAB16A* transcripts were induced even at 2 d and continued to increase for 6 d (Fig. 3B). *OsPT11* mRNA levels did not significantly decline in response to WD until 6 d but partially recovered one day after rewatering. After 8 d WD, recovery of *OsPT11* expression was severely attenuated across all samples. When the recovery period was extended to 28 d we found that *OsPT11* expression not only recovered but surpassed WW control plants (Fig. 3B). These results demonstrate AM colonization is highly dynamic in response to external environmental stimuli, resulting in transitory quiescence and resurgence in symbiotic status, both influenced by the duration and extent of the stress.

Finally, we sought to determine whether similar observations are evident in the field where soil composition is more complex and plant water deficit progresses more slowly than in pot-grown plants, due to buffering by the water table. After transplantation into a waterlogged paddy and an initial 2 week acclimation period, the plot was drained by gravity and plant root samples were collected weekly. The symbiosis marker transcripts *OsAM1* and *OsPT11* were very low as the paddy drained and gradually increased over 6 weeks, at which point irrigation was stopped. *OsRAB16A* transcripts increased within a week of drying and were stably high for 4 weeks, at which point leaf rolling was observed and the plot was irrigated. *OsAM1* and *OsPT11* transcript levels remained stable until week 8 when levels fell precipitously. After a 1 week re-irrigation period, these transcripts slightly increased but were not significantly elevated (Fig. 3C). These results validate pot demonstrations that AM symbiosis is highly dynamic to non-nutritional environmental stressors.

## Discussion

The intricate AM-plant interaction simultaneously confers benefits and costs to both the host and the symbiont. The plant bears considerable cost in terms of carbon provision to the symbiont, a fatty acid heterotroph. However, this reliance on lipid provision may also act as an important lever for modulating fungal activity under diverse conditions. During drought, photosynthates become limited and in the symbiotic economy are increasingly scarce. The plant prioritizes valuable resources for root pioneering for moisture^23–25^ and investment in lipid barriers to limit water loss^7^. Yet, counterbalancing the loss in host carbon fixation may be the provision by AM of mineral nutrients in an aqueous stream from hyphae that extend beyond the root zone^26^. Our study shows that when water becomes limiting the plant host disengages with the symbiont, conserving fixed carbon for its own means, a phenomenon that appears conserved in both rice and tomato.

Fungal retraction of septae and maintenance of vesicles, which are robust propagules^27^, appears to safeguard AM, allowing for *de novo* formation of active arbuscles upon reirrigation. AMF lifespan is independent from its host despite their obligate biotrophy. AMF propagules out-survive their hosts^27^ and can continue to grow and acquire nutrients from dead host roots^28,29^. Mycelial growth and even arbuscular viability is detectable for several months after photosynthate delivery ceases due to shoot detachment from host chicory plants^30^, suggesting that SDH-stained arbuscules that are lost during drought is not solely due to reduction in photosynthesis. The reduction in plant nutrient acquisition programs points to a broader reduction in nutrient demands which could have significant impacts on shaping microbial communities. The balance between photosynthetic reduction and plant-derived signaling, *e*.*g*., repression of PSR genes (Fig 1C), in maintaining intraradical viability during water deficit deserves further consideration. This is relevant to optimizing the benefits of alternating wet and dry cultivation of rice, a promising approach for reducing overall methane emissions from anaerobic methanogenic bacteria and reducing freshwater consumption of traditional paddy cultivation^7,8^.

The need for dynamic flexibility in modulating symbiotic status highlights potential challenges associated with engineering AM symbiosis and other mutualistic interactions in field crops. A promising strategy to improve AM benefit is to engineer phosphate-insensitive symbiosis^31^; however, doing so could confer insensitivity to external stimuli such as drought. We hypothesize that a sustained symbiotic program during drought would *reduce* plant resiliency to WD as the symbiont would be a carbon sink that diverts resources from the plant’s own drought resilience strategy. Thus, a nuanced approach to tuning symbiosis may be needed for crops to retain dynamism to other stressors.

## Materials and Methods

### Plant materials, growth and colonization

*Oryza sativa* cv. Nipponbare were surface sterilized in 50% (v/v) bleach for 10 min and germinated in sterile water for 3 d in the dark at 30 °C. 5 seedlings were transplanted into pots of sand and thoroughly washed Profile Greens Grade mixed in a ratio of 7:3. For mycorrhizal colonization, seedlings were instead transplanted into a mixture of 13:6:1 sand:Profile Greens Grad®:Mykos Gold Granular *Rhizophagus irregularis* inoculum (Reforestation Technologies International). Plants were grown in a glasshouse in Riverside, CA maintained with a daytime temperature of 30 °C and nighttime temperature of 25 °C. Plants were drip-irrigated with low P fertilizer as described^32^, pots were allowed to drain to permit oxygenation to support fungal growth. For experiments using *smax1* mutants^33^, the corresponding wild type background Dongjin was used as a control and were grown as described for Nipponbare.

Rice water deficit was initiated 6 wk post-seeding when plants had developed two tillers. Watering was stopped and samples were collected at solar noon the day that leaf rolling was observed in the morning, after 5-6 d. To examine WD recovery, pots were re-watered until the growth substrate was fully saturated; samples were harvested after 24 h. For collection, plants were released from the pots, excess growth substrate was removed by gentle shaking and roots were cut at the base of the crown. To separate deep and shallow roots, root systems were cut 7.5 cm below the crown. To collect crowns, any root remnant was carefully removed and a 1 cm portion measured up from the base of the crown was saved for future analysis.

For tomato experiments, *Solanum lycopersicum* M82 seeds were imbibed in water on paper towels for 3 d in the dark at room temperature. Granular inocula of *Rhizophagus irregularis* (DAOM197198, MYKOS GOLD WP, RTI, Gilroy, CA) or Field & Fairway (Profile, non-Myc control) were sprinkled on those germinated seeds and kept at room temperature for 1 d. Seedlings were transferred to custom-made black 50 mL growth tubes; the bottom two-thirds was filled with sand while the top one third was filled with with AMF/sand mix (1:3 vol, Myc plants) or Field Fairway/sand mix (1:3 vol, Non-myc plants). Drainage was by four holes drilled into the bottom of each growth tube. Plants were irrigated with 1 mL of modified half strength Hoagland’s solution^34^ containing 25 μM phosphate at the following frequencies: week 1-3, 3 times per week; week 4-5, 2 times per week; week 6, 4 times per week. Plants were covered with a clear humidity dome; the dome vent was opened at week 5.

Tomato water deficit was initiated after 6 wks of growth. WD-treated plants had irrigation withheld while WW-treated plants were irrigated daily. Plants were removed from the tubes and roots were cut 1 cm below the substrate surface. The roots of 6-8 plants were pooled and frozen in liquid nitrogen.

### RNA and DNA extraction, sequencing and quantitative PCR

For rice, flash frozen tissue was ground in a mortar and pestle under liquid nitrogen. mRNA isolation was performed and RNA sequencing libraries were prepare exactly as described previously^7^ and were sequenced with either Illumina NextSeq 550 at the UCR Genomics Core or NovaSeq at Novogene. For quantitative RT-PCR (qPCR), isolated polyA mRNA was reverse transcribed with Maxima RT (ThermoFisher Scientific) and the resulting cDNA was diluted 10-fold for qPCR with SsoAdvanced Universal SYBR Green Supermix (BioRad) on a BioRad CFX-384 Real-Time PCR instrument following the manufacturer’s instructions using the primers listed in Table S13.

For tomato, samples were ground in a bead beater and mixed with 600 µl Trizol. RNA was extracted using the Direct-zol RNA Microprep Kit (Zymo Research, Irvine, CA, USA). Poly-A-primed libraries were constructed and sequenced by Novogene.

### RNA seq data processing and analysis

Transcriptome data was trimmed with cutadapt. Rice root samples were sequenced with either 75 bp or 150 bp reads; 150 bp reads were trimmed to 75 bp to be consistent across samples. For data alignment, the transcriptomes of *R. irregularis* and either host plant were concatenated and used as an input for pseudoalignment with kallisto^35^. Transcriptome annotations for *R. irregularis* were from Manley *et al*^*12*^. For tomato, ITAG 4.0 annotations were obtained from Sol Genomics^36^. For rice, RAP-DB^37^ annotations were obtained from Ensembl Release 56^38^. Gene level summarization was performed using tximport^39^. Differentially expressed genes were called with DESeq2^40^. DEseq2 analysis was performed separately for plant and fungal genes. For plant genes, the DEseq2 design matrix included both colonization (Myc and Non-myc) and water status (WW and WD), simultaneously. Co-expression networks were created as described previously by Wisecaver *et al*.^*41*^. Coexpression analysis was performed using the combined rice/AMF dataset using only colonized Myc samples. Co-expression networks were made using decay constants of 5, 10, 25, 50, 100 and 200 and an adjusted *p* value of 0.1 was used as a significance cutoff.

GO term enrichment and visualization was performed with cluster profiler^42^. Rice GO terms were curated by combining GO terms from RAP-DB, Oryzabase and GO terms for Arabidopsis orthologs. To identify Arabidopsis orthologs, 1:1 associations were determined by OrthoFinder^43^ and corresponding GO terms were obtained from the Araport 11 annotations^44^. For *R. irregularis*, GO terms were curated by compiling GO terms assigned by Manley et al^12^ and GO terms from orthologous *Saccharomyces cerevisiae* genes which were identified in the same manner as described for rice-Arabidopsis orthologs. GO term annotation files were then constructed as described previously^45^.

## Phylogenetics

Phylogenetic tree construction was performed as described previously^7,46^. Fungal species used for phylogeny construction are listed in Table S11. Orthogroups were created with OrthoFinder^43^ using species listed in Table S11.

## WGA and SDH staining

For WGA staining, root systems were fixed in 80% (v/v) EtOH, cleared in 10% (w/v) KOH for 30 min at 90 °C, briefly rinsed in 1% (w/v) HCl, rinsed 3 times in water and finally washed in Tris-buffered saline (TBS), pH 7.4. Root samples were stained overnight in 1 μg/ml WGA-FITC (VectorLabs) in TBS overnight. Colonization frequency was assessed on a Leica microscope equipped with a Chroma 41001 EGFP filter cube using the gridline intersect method^47^.

For SDH staining, roots were harvested into TBS and stained for SDH overnight as described^48^. Roots were rinsed in water and cleared in ClearSee^49^ for several days. Colonization frequency was determined with a light microscope using the gridline intersect method.

To quantify septation, WGA-stained root sections were imaged on a Zeiss 880 confocal microscope in AiryScan mode with an Argon laser (Zeiss) at 488 nm excitation. Imaging was performed with a Plan-Apochromat 10x/0.45 objective and 2x digital zoom. To ensure that comparable root segments were used for analysis, large lateral root segments containing fully developed arbuscules were chosen for imaging. Z stacks were collected for each root and Airyscan processing was performed with the auto settings in the Zeiss Zen software. For quantification, the maximum projection image was created in ImageJ. The total length of all longitudinal hyphae was measured using the segmented line tool in ImageJ and the septa along the measured hyphae were counted using the multi-point tool.

### Inductively coupled plasma-optical emission spectroscopy (ICP-OES)

For elemental analysis, frozen shoot base tissue (1 cm as measured from the shoot base) was pulverized and an aliquot of tissue was dried at 65 °C for 3 days. The tissue was digested in 1 ml 70% (v/v) trace metal grade nitric acid for 30 min at 95 °C; 0.5 ml hydrogen peroxide was then added and digested again for an additional 30 min at 95 °C. Samples were centrifuged at 3220 *g* for 10 min to pellet insoluble material and diluted 10-fold in water before analysis. ICP-OES was performed with a Perkin-Elmer Optima 7300DV at the Earth Sciences Research Laboratory at UC Riverside.

## Data Accessibility

All sequencing reads are available through NCBI GEO (GSE304369).

We thank Gabriel Ferreras Garucho and Tania Chancellor for helpful suggestions. Research by the authors is supported by the US National Science Foundation IOS-211980 (to J.B.S.) and DGE-1922642 (to J.B.S. and A.T.B) and the US Department of Agriculture, US National Institute of Food and Agriculture - Agriculture and Food Research Initiative Grant no. 2019-67013-29313 (to J.B.S.), no. 2022-67012-36716 (to G.Z.A) and no.1026477 (to A.T.B.).

